# Apigenin Improves Healthspan and Protects Against Alzheimer’s Disease-Associated Proteotoxicity in *C. elegans*

**DOI:** 10.64898/2026.09.23.753917

**Authors:** Randy A. Grant, Lauren H. E. Park, Devin Wahl, Thomas J. LaRocca

## Abstract

Apigenin is an abundant, bioactive phytochemical with reported therapeutic potential and neuroprotective effects in a variety of age-related disease models. However, the extent to which apigenin may promote healthspan and protect against neurodegeneration-associated proteotoxicity has not been thoroughly investigated, and the biological processes/mechanisms that underlie its protective effects remain unclear. To determine if apigenin increases healthspan, we performed lifespan assays in conjunction with measures of aging-relevant physiological function in wild-type *C. elegans*. We found that chronic exposure to apigenin extended median lifespan and improved multiple readouts of healthspan, including pharyngeal pumping, locomotor activity, and age-related macromolecular damage accumulation. Additional analyses of transgenic reporter strains and transcriptomic profiling (RNA-seq) indicated that apigenin primarily modulated redox-sensitive stress signaling and pathways associated with cellular homeostasis in aging animals. Finally, in transgenic *C. elegans* strains expressing the pathological Alzheimer’s disease-associated proteins amyloid-beta or tau, we found that apigenin had similar beneficial effects on lifespan, healthspan, and proteotoxicity-associated functional impairments. Together, our results support apigenin as a potential healthspan-promoting intervention that acts in part through reductions in oxidative stress-associated signaling, with beneficial effects that extend to models of Alzheimer’s disease-associated proteotoxicity.

## INTRODUCTION

Aging is characterized by progressive declines in multiple cellular and physiological functions that increase susceptibility to chronic diseases, including neurodegenerative disorders (1–5). Therefore, as the global population ages (6), there is a growing need for interventions that increase healthspan and reduce age- and disease-related pathological events (7). In this context, phytochemicals (naturally occurring bioactive dietary compounds with health-promoting and therapeutic effects) represent one promising class of interventions (8–10). Recently, the phytochemical apigenin, found naturally in chamomile and parsley, has gained attention for its reported therapeutic effects in various age-related disease models (11–14). However, the exact molecular mechanisms underlying these broad therapeutic effects are incompletely understood.

We recently reported that apigenin crosses the blood-brain barrier and exerts beneficial effects of apigenin in the aging mouse brain (11), and other data from model organisms suggest apigenin may attenuate pathology associated with Alzheimer’s disease (AD) (15–17). Brain aging is tightly linked with the development of neurodegenerative diseases like AD (18), and many older, disease-free adults exhibit pathological AD protein accumulation (e.g., amyloid-beta [Aβ] and tau). As such, and because there remain no effective treatments for preventing or reversing brain aging/AD (19–23), identifying safe, long-term interventions that preserve neurological healthspan is an important research direction. Interestingly, apigenin has been reported to modulate inflammatory signaling, antioxidant function, cell cycle regulation, and apoptosis pathways (11,24–26). However, like many phytochemicals, it has also been proposed to exert its beneficial effects through hormetic mechanisms involving transient activation of cellular stress response pathways. Yet, exactly which pathways/mechanisms are activated by apigenin in the context of aging and AD-associated proteotoxicity remains unclear.

In the present study, we evaluated apigenin’s effects on lifespan, age-related declines in physiological function, and the adverse consequences of AD-associated proteins in *C. elegans*, while using RNA-seq and *in vivo* fluorescent reporter strain imaging to characterize underlying molecular mechanisms. We show that apigenin extends median lifespan in wild-type animals as well as transgenic AD proteinopathy models, and that these effects are associated with improved neuromuscular function and reduced age-related macromolecular damage accumulation. Our RNA-seq and imaging data also identify reductions in basal oxidative stress-associated signaling as a key consequence underlying the therapeutic effects of apigenin. Together, our results add further support to the growing data on apigenin as a promising candidate for preserving healthspan and, perhaps, delaying neurodegeneration.

## METHODS

### *C. elegans* Strains, Maintenance, and Treatment

Wild-type and transgenic *C. elegans* strains used in this study (Table 1) were purchased from the Caenorhabditis Genetics Center at the University of Minnesota and maintained on standard nematode growth medium (NGM) plates containing _CaCl2_, MgSO_4_, KPO_4_, and 5 mg/mL cholesterol (27). For treatments, animals were plated with or without 50 µM apigenin (Cayman Chemical), which was supplemented into the *E. coli* OP50 food source, and housed at 20 °C unless otherwise stated. This dose was selected to model relevant physiological exposure levels achievable through dietary or nutraceutical intake in humans. Previous studies have demonstrated that daily consumption of apigenin-rich foods results in a human plasma concentration of approximately 0.13 µM (28), while semi-purified nutraceutical capsule intake can raise circulating levels to 1-5 µM (29). In *C. elegans* models of aging, oral exposure to compounds present in the food source typically yields steady-state tissue concentrations of 1-4% of the dose plated (30). Therefore, the concentration of 50 µM used in this study is estimated to achieve tissue concentrations of 0.5-2.0 µM, which are comparable to feasible, non-toxic *in vivo* human circulating levels following dietary or supplemental consumption. For experiments that involved aging worms, NGM plates were supplemented with 400 µM 5-Fluoro-2’-deoxyuridine (FUdR, Research Products International) directly into the molten agar to maintain age-synchronized populations. FUdR was not used for lifespan experiments, and for all other experiments, control and apigenin-treated animals were exposed to identical FUdR concentrations to minimize potential confounding effects.

**Table 1.** *C. elegans* Strains used in the present study. The table details specific strain designations, phenotypic descriptions of the fluorescent reporters or disease models, and complete genetic configurations.

| Strain | Description | Genotype |
| --- | --- | --- |
| LD1171 | GFP reporter for *gcs-1* expression | ldIs3 [*gcs-1p*::GFP + *rol-6*(su1006)] |
| LD1 | GFP reporter for *skn-1b/c* expression | ldIs7 [*skn-1b/c*::GFP + *rol-6*(su1006)] |
| OH13908 | mKate reporter for *daf-16* expression | *daf-16*(ot821[*daf-16*::mKate2::3xFLAG]) |
| SJ4103 | GFP-labeled muscle mitochondria | zcls14 [*myo-3*::GFP(mit)] |
| IG274 | GFP reporter for *nlp-29* expression | frIs7 [*nlp-29p*::GFP + *col-12p*::DsRed] |
| DV3285 | mKate reporter for *mpk-1* expression | *his-72*(cp76[mNeonGreen::3xFlag::*his-72*]) *mpk-1*(re172[*mpk-1*::mKate2::3xFlag]) |
| GRU102 | Pan-neuronal human Aβ<sub>1-42</sub> expression | gnals2 [*myo-2p*::YFP + *unc-119p*::Abeta1-42] |
| BR5706 | Pan-neuronal pathological human tau expression | byIs193 [*rab-3p*::F3(delta)K280 + *myo-2p*::mCherry].<br>bkIs10 [*aex-3p*::hTau V337M + *myo-2p*::GFP] |
| CL4176 | Temperature-inducible muscle Aβ expression | dvIs27 [*myo-3p*::A-Beta (1-42)::let-851 3'UTR) + *rol-6*(su1006)] |
| KAE112 | Muscle tau expression | seals201 [*myo-3p*::human tau (0N4R;V337M)::unc-54 3'UTR + *vha-6p*::mCherry::unc-54 3'UTR] |
| NC1686 | GFP-labeled PVD neurons | wdIs51 [F49H12.4::GFP + *unc-119*(+)] |

### Synchronization and Lifespan Assays

Animals were grown on NGM plates and then age-synchronized by isolating eggs via standard sodium hypochlorite protocols (31). Briefly, worm populations were washed and collected with M9 buffer and centrifuged at 1,700 RPM for 1 min. The pellet was treated with ∼2% sodium hypochlorite followed by 0.4 M sodium hydroxide. The solution was vortexed and centrifuged again, and the pellet was resuspended in sterile water, then centrifuged and washed with sterile water three times before being resuspended in sterile M9 buffer. Eggs were left to hatch in M9 buffer for 18 to 24 hours before being distributed on NGM plates and allowed to mature. Lifespan experiments were performed using a Nemalife Infinity apparatus (NemaLife Inc.) as previously reported and according to manufacturer’s instructions (32,33). Briefly, 100% ethanol was used to clean Nemalife microfluidic chips and then displaced with 50 mg/mL pluronic F-127 (Sigma-Aldrich). Chips were incubated at room temperature for 1 hour and then thoroughly washed with M9 buffer. Then, 50 – 75 untreated day 1 *C. elegans* adults were added to each chip. All chips were maintained at 20 °C and were washed, fed/treated (20 mg OP50/mL NGM solution), and video recorded daily. Videos were used to quantify living animals each day (washing/feeding stimulates nematode movement), and animals were scored as alive if movement was observed during video duration.

### Pharyngeal Pumping

Pharyngeal pumping was quantified as reported elsewhere (34) at estimated median lifespans (N2: day 10, GRU102 & BR5706: day 7). Individual *C. elegans* pharynges were observed in Nemalife chips placed under a stereo microscope (Tritech Research) at 50x magnification. Contractions were manually recorded over a 60-second interval for 5 to 10 animals per condition.

### Velocity Analysis

All velocity data were gathered using WormLab (MBF Bioscience LLC) software (35) from lifespan videos (collected as described above) or in videos of KAE112 paralysis experiment plates. Briefly, all videos were trimmed to 30 seconds and cropped to include only the microfluidic arena or OP50 lawn on plates. In WormLab, the arena was measured to calculate pixel size of the resized video, the “light worms on a dark background” setting was selected, and the threshold level was adjusted for optimal contrast between the worms and their background. Tracking was analyzed using default parameters except for the following changes: ‘detection parameters’ – area min: 0, length min: 1, width min: 0.5, width/length ratio min: 0.05, max: 0.3, detection fit min: 0.25, registration fit: 0.05, detection frequency: 15 frames, predominant direction: forward, fitting iterations: 130; ‘tracking parameters’ – max tracked hypothesis: 3, minimum track duration: 3. After WormLab’s automated tracking, individual worm tracks were manually repaired using the “remove” and “join” track functions. Moving average speed was exported to Excel with maximum and average velocities calculated using standard functions. Only velocity data for worms tracked for more than half of the video duration were included in the final analyses to eliminate possible repeated sampling of the same worm.

### Imaging

Live animals were placed on a thin 5% agar pad in water, immobilized in 3% 1-phenoxy-2-propanol (Tokyo Chemical Industry) in water, and imaged using an EVOS M7000 fluorescence microscope (ThermoFisher) with DAPI (Ex; 357, Em; 447), GFP (Ex; 470, Em; 525), and RFP (Ex; 531, Em; 593) LED light cubes. NC1686 worm strain neurodegeneration images were captured with an Olympus UplanSApo 40x/0.95 NA objective, and all other images were captured with an Evos Plan FL PH 10x/0.30 NA objective. For neurodegeneration imaging, a ∼20-plane Z-stack was captured, and all images were deconvolved using Celleste (ThermoFisher) imaging software before analyzing dendrite degeneration. Briefly, contrast was increased equally across all images for visualization and quantification of quaternary (4°) processes. 4° processes were defined as terminal perpendicular dendritic branches forming the outermost structures of the PVD menorahs. Images were then reset to the deconvolved output for manual scoring of dendritic blebs (a marker of degeneration of neurons). All neurodegeneration data were normalized to primary PVD dendrite length with quantifications compared against a representative pool of day 1 baseline animals that were split into treated and untreated experimental groups. Fluorescence intensity of age-related autofluorescence and reporter strains was quantitatively analyzed in FIJI (NIH). Briefly, light intensity was kept consistent throughout entire experiments as whole worms were imaged. When quantifying fluorescent intensity in FIJI, whole worms were defined as the region of interest and standard protocols for calculating corrected total fluorescence (which includes background subtraction) were used (36).

### RNA Extraction/Bioinformatics

Wild-type *C. elegans* with or without apigenin treatment were aged to day 5 or 10 (to approximate younger vs. late middle life when healthspan/lifespan effects were strongest). Triplicates of ∼100 animals for each condition were collected in standard M9 buffer. Trizol Reagent (Zymo Research) was mixed into each sample, and samples were subsequently frozen in liquid nitrogen, thawed at room temperature, and vortexed three times before being centrifuged at 12,000 RPM for 30 seconds. Supernatant was carefully transferred to a new microfuge tube and 100% ethanol was added. RNA was recovered and purified using an RNA-specific spin column kit (Direct-Zol, Zymo Research), which included DnaseI treatment to remove genomic DNA. Poly-A RNA libraries were generated using a Tecan NuGEN library kit, with libraries sequenced on an Illumina NovaSeq 6000 platform to generate >40 M 150-bp paired-end reads per sample (Genomics Core, University of Colorado Anschutz Medical Campus).

Transcriptome data analyses were performed as recently reported (11,32,37–40). Reads were trimmed and filtered with the *fastp* program (41), then mapped to the *C. elegans* (ce11) genome with the STAR aligner (42). Gene counts were analyzed for differential expression in R using DESeq2 software (43), with statistically significant differences determined using the default Wald test with *p*-values adjusted for multiple testing using the Benjamini-Hochberg false discovery rate (FDR) correction. Normalized counts extracted from DESeq2 were used for all statistical analyses, and gene ontology (GO) was performed using g:Profiler (44).

### Paralysis Assays

Paralysis experiments for Aβ-expressing strains were performed as reported elsewhere (45). Briefly, untreated synchronized *C. elegans* in their third larval stage (L3) were transferred to new NGM plates with or without treatment. These new plates were transferred from 16 to 25 °C and incubated for 15 hours. After the incubation period, *C. elegans* were assessed for paralysis every two hours until all animals were completely paralyzed. Total paralysis was scored when animals were prodded with a pick and no movement was observed. Tauopathy strain paralysis experiments were performed similarly and as reported elsewhere (46). Briefly, untreated synchronized *C. elegans* in their fourth larval stage (L4) were transferred to FUdR plates with or without treatment. On days 2-10 of adulthood, 5-min video recordings were taken with Nemalife imaging equipment and velocity analysis was performed in WormLab as described above in section 2.4.

### Statistical Analyses

Statistical analyses for lifespan assays were performed using a log-rank (Mantel-Cox) test. For all other data, normality was assessed using the Shapiro-Wilk test, and comparisons between treated and untreated animals were performed using two-tailed Student’s *t*-tests. For neurodegeneration analyses involving multiple groups, one-way ANOVA followed by multiple comparisons correction was used. All statistical analyses were performed using GraphPad Prism, except for RNA-seq data (described above under section 2.6), for which differential expression analysis was performed using DESeq2 software with significant differences determined using the default Wald test, and in which GO significance was determined using the R package g:Profiler-derived *p*-values.

## RESULTS

Apigenin has been reported to have therapeutic effects in multiple settings/models of disease (47), including brain aging (11), yet the extent to which it may protect against healthspan and neurodegeneration, and the mechanisms involved, have not been thoroughly investigated. To provide initial insight into the efficacy and mechanisms of action for apigenin in this context, we first performed lifespan assays on treated and untreated wild-type (N2) *C. elegans* in microfluidic chips, as previously reported (32,33). We observed a modest but significant extension in median lifespan for worms exposed to apigenin (**Fig. 1a**), and a slight but non-significant increase in maximum lifespan. In the same animals, neuromuscular function (a key metric of healthspan (48,49)) was improved at day 10 (“late middle age”) in treated animals, as reflected by significant increases in pharyngeal pumping and both average and maximum crawling velocity (**Fig. 1b,c**). We also measured whole worm 447 nm autofluorescence, an indicator of oxidized/cross-linked macromolecule and gut granule accumulation that has been linked to age-related declines in *C. elegans* (50), and we found that autofluorescence was significantly lower in day 10 apigenin-treated worms compared to controls **(Fig. 1d,e**). Together, these data suggest that apigenin attenuates age-related functional declines (i.e., improves healthspan) in wild-type *C. elegans*.

**Fig. 1.**
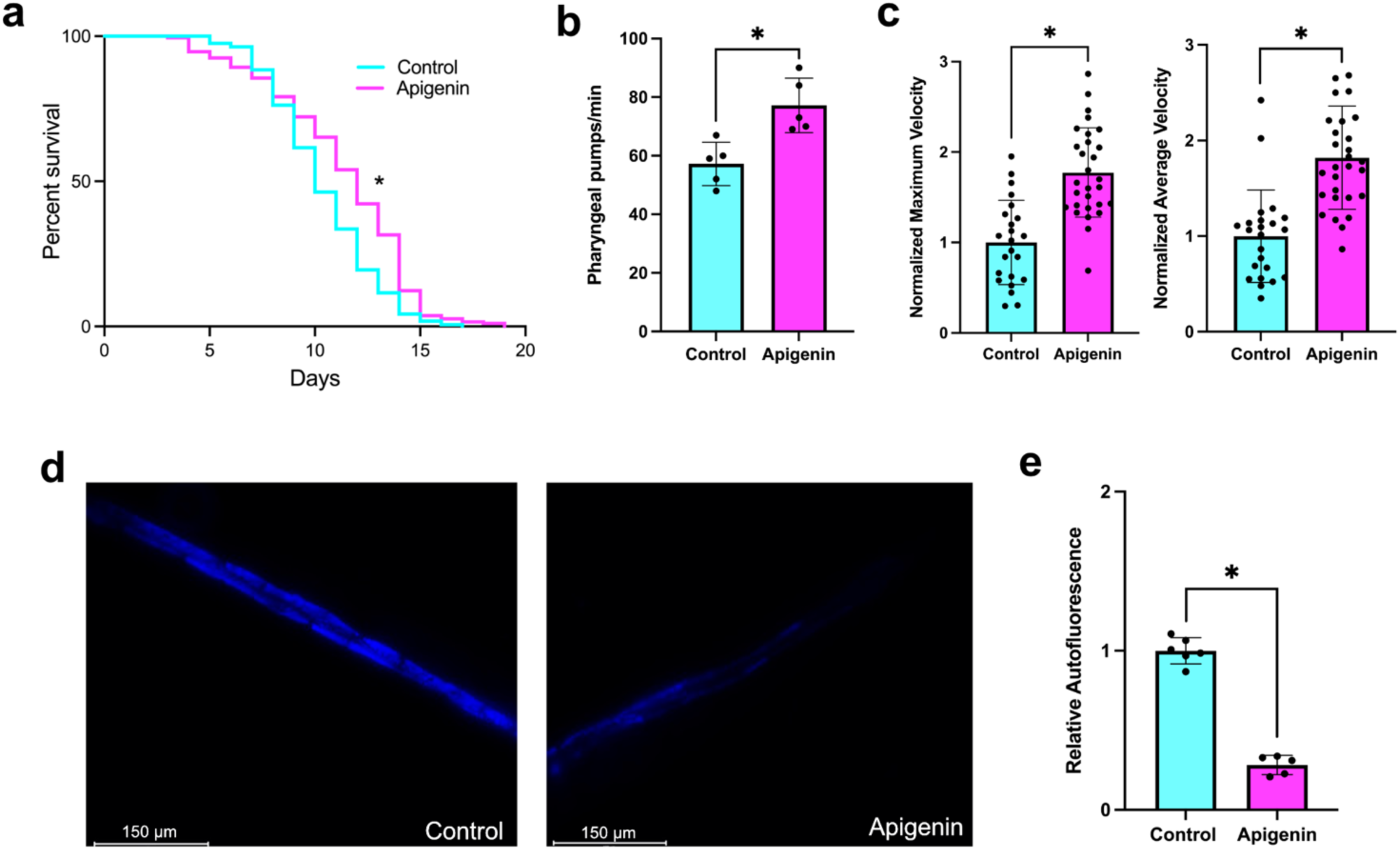
Apigenin attenuates age-related functional declines in wild-type *C. elegans*. **a)** Median lifespan was significantly extended in apigenin-treated worms (control n = 164, apigenin n = 187). In the same animals, neuromuscular function was significantly enhanced with treatment, as reflected by increased **b)** pharyngeal pumping rates (*n* = 5/group) and **c)** average and maximum crawling velocity (*n* = 20-30/group). **d)** Representative age-related autofluorescence (λ = 447 nm) images (*n* = 5-6/group). **e)** Quantification of age-related autofluorescence intensity. All measurements were assessed at Day 10. \**p* < 0.01

To examine the effects of chronic apigenin exposure on age-related neurodegeneration, we imaged sensory PVD neurons *in vivo* in treated and untreated *C. elegans* expressing a GFP reporter at three timepoints (**Fig. 2a**) as previously reported (32,51,52). PVD neurons are commonly analyzed in neurodegeneration studies for their extensive dendritic arborization and ease of visualization (53), with the quantification of spherical dendritic “blebs” serving as a well-characterized marker of neuronal degeneration (54). While overall dendritic architecture was preserved in both control and treated animals (**Fig. 2b**), we observed a significant age-associated accumulation of neuronal blebbing in control worms, which did not reach significance in apigenin-treated animals (**Fig. 2c**). However, direct comparisons between control and apigenin treated cohorts showed no significant differences at individual timepoints, suggesting that while apigenin may mitigate age-associated changes in neuronal structure *in vivo*, its overall effects may be modest.

**Fig. 2.**
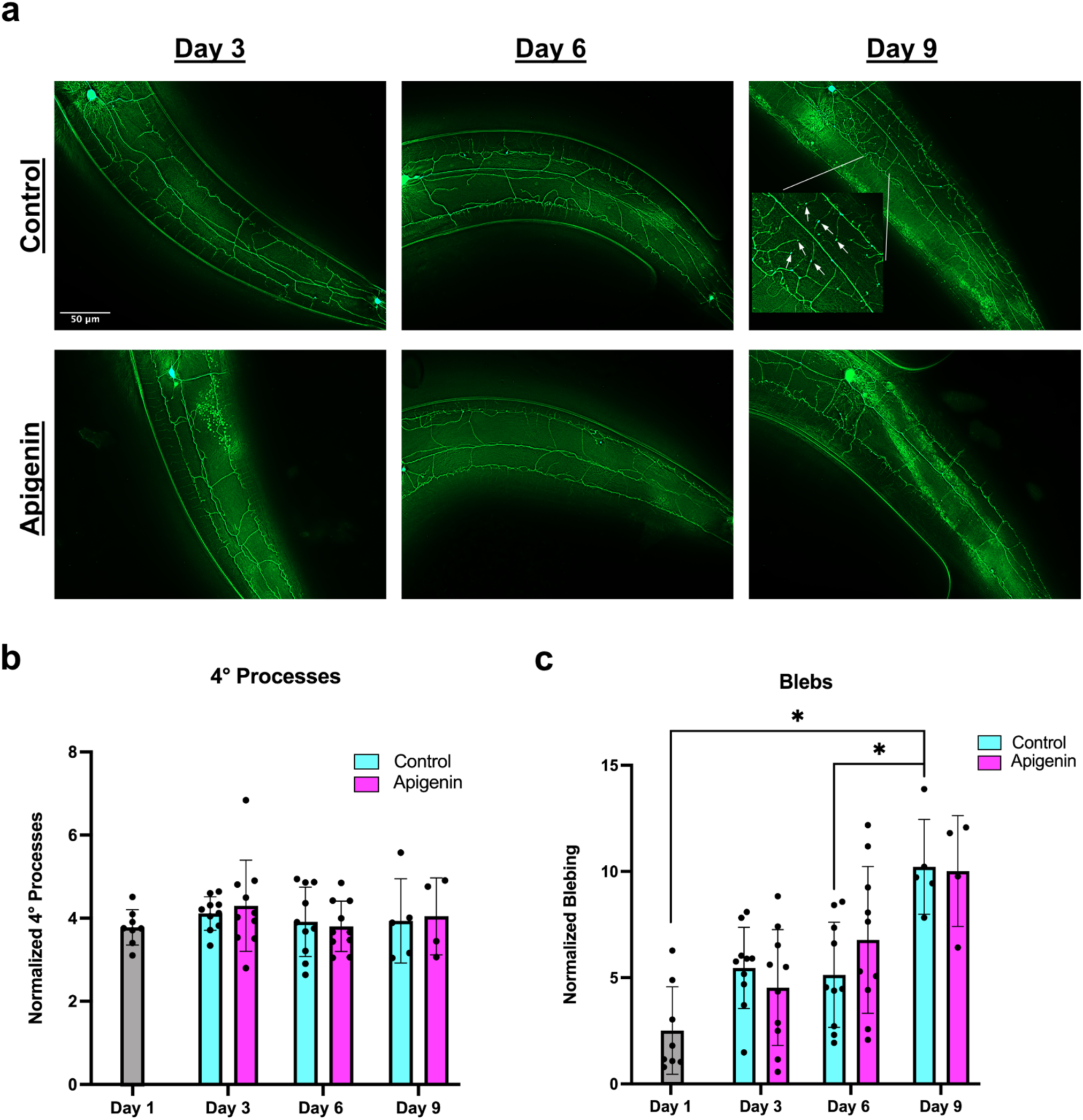
Effects of apigenin on age-related PVD neurodegeneration. **a)** Representative PVD fluorescence images of control and apigenin-treated NC1686 *C. elegans* compared against day 1 baseline animals. Quantification (*n* = 4-10/group) of **b)** quaternary processes and; **c)** dendritic blebbing normalized to primary dendrite length. \**p* < 0.05

Although apigenin has been shown to be protective in a variety of age-related disease models (26,47,55), reports on the primary biological processes affected by apigenin in these settings are mixed. Therefore, to better understand the mechanisms underlying apigenin’s therapeutic effects and our observations above, we performed *in vivo* fluorescence imaging on various day 10 adult transgenic *C. elegans* reporter strains with and without chronic apigenin treatment. Interestingly, apigenin-treated worms showed significantly less expression of proteins involved in two key *C. elegans* oxidative stress pathways, glutamate-cysteine ligase (*gcs-1p*) and skinhead-1 (*skn-1b/c*) (**Fig. 3a,b**). Consistent with a state of reduced systemic stress, expression of *daf-16*, a FoxO ortholog tightly associated with stress responses and biological aging (56), was also reduced in treated animals (**Fig. 3c**). Despite these observations, we did not observe any significant differences in gross mitochondrial-associated GFP signal (**Fig. 3d**) or markers of innate immune activation (**Fig. 3e**). However, the MAPK pathway (as reflected by *mpk-1*), which is centrally involved in both oxidative stress and immune signaling (57,58), was significantly reduced with apigenin treatment (**Fig. 3f**).

**Fig. 3.**
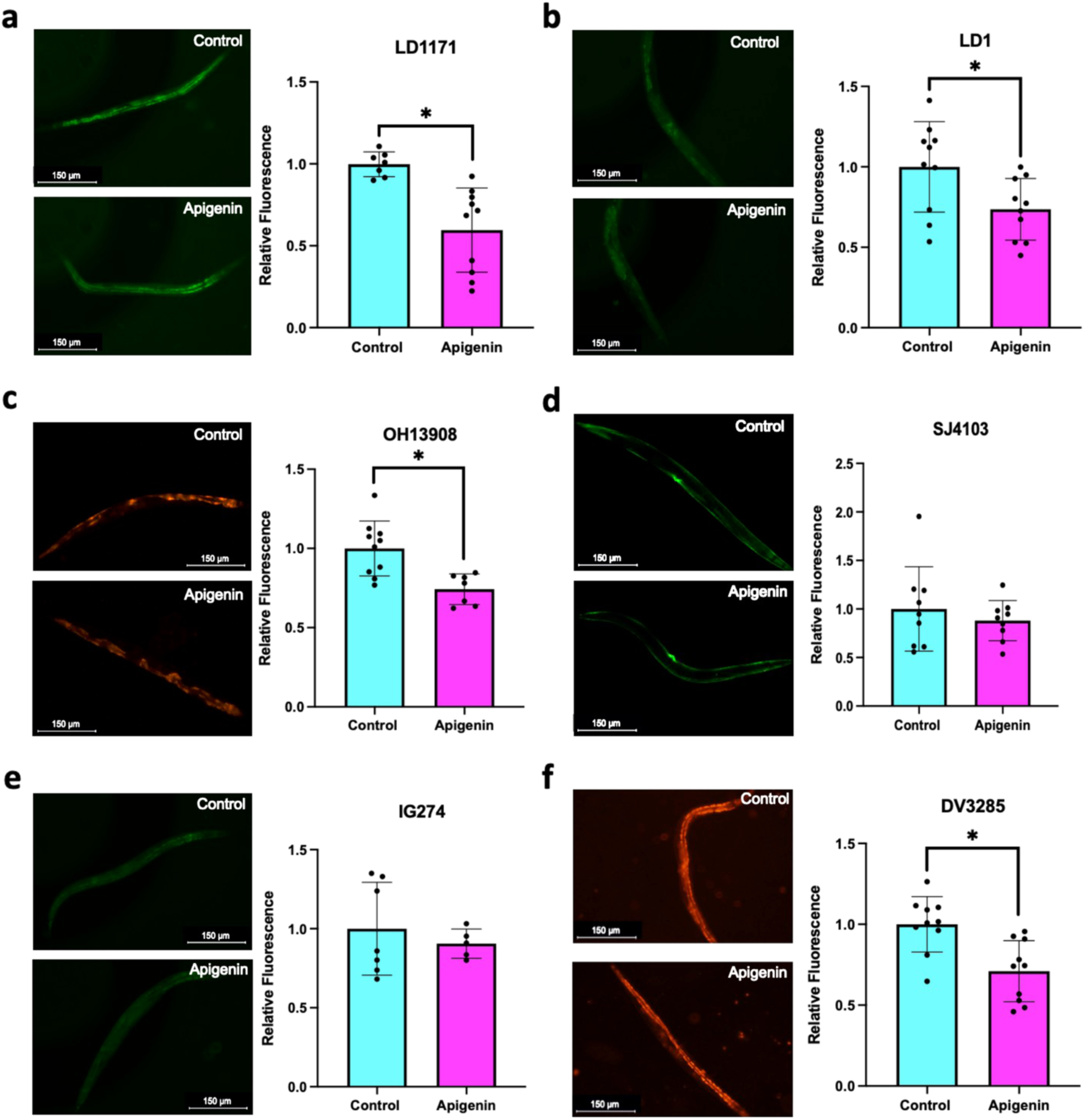
Apigenin reduces oxidative stress-associated signaling with aging. Effects of apigenin treatment in transgenic *C. elegans* strains expressing fluorescent reporters for: **a)** glutamate-cysteine ligase (*gcs-1p*::GFP); **b)** oxidative stress resistance (*skn-1b/c*::GFP); **c)** age- and stress-related insulin/IGF-1 signaling (*daf-16*::mKate2); **d)** muscle mitochondria (*myo-3*::GFP); **e)** innate immune activation (*nlp-29p*::GFP); and **f)** MAP kinase expression (*mpk-1*::mKate2). All images were captured at day 10 (*n* = 5-10/group). \**p* < 0.05

Together, these data suggest apigenin reduces the activation of endogenous age-associated stress response pathways, and especially oxidative stress signaling networks.

To characterize wider biological changes induced by chronic apigenin exposure during aging, we performed RNA-seq on control and apigenin-treated *C. elegans* at day 5 and day 10. In differential gene expression analyses, we identified 896 significantly different transcripts in old vs. younger control animals, compared to 703 significantly different transcripts in old vs. younger treated animals (**Fig. 4a,b**), potentially suggesting a modest suppression of age-related transcriptome changes in apigenin-treated animals.

**Fig. 4.**
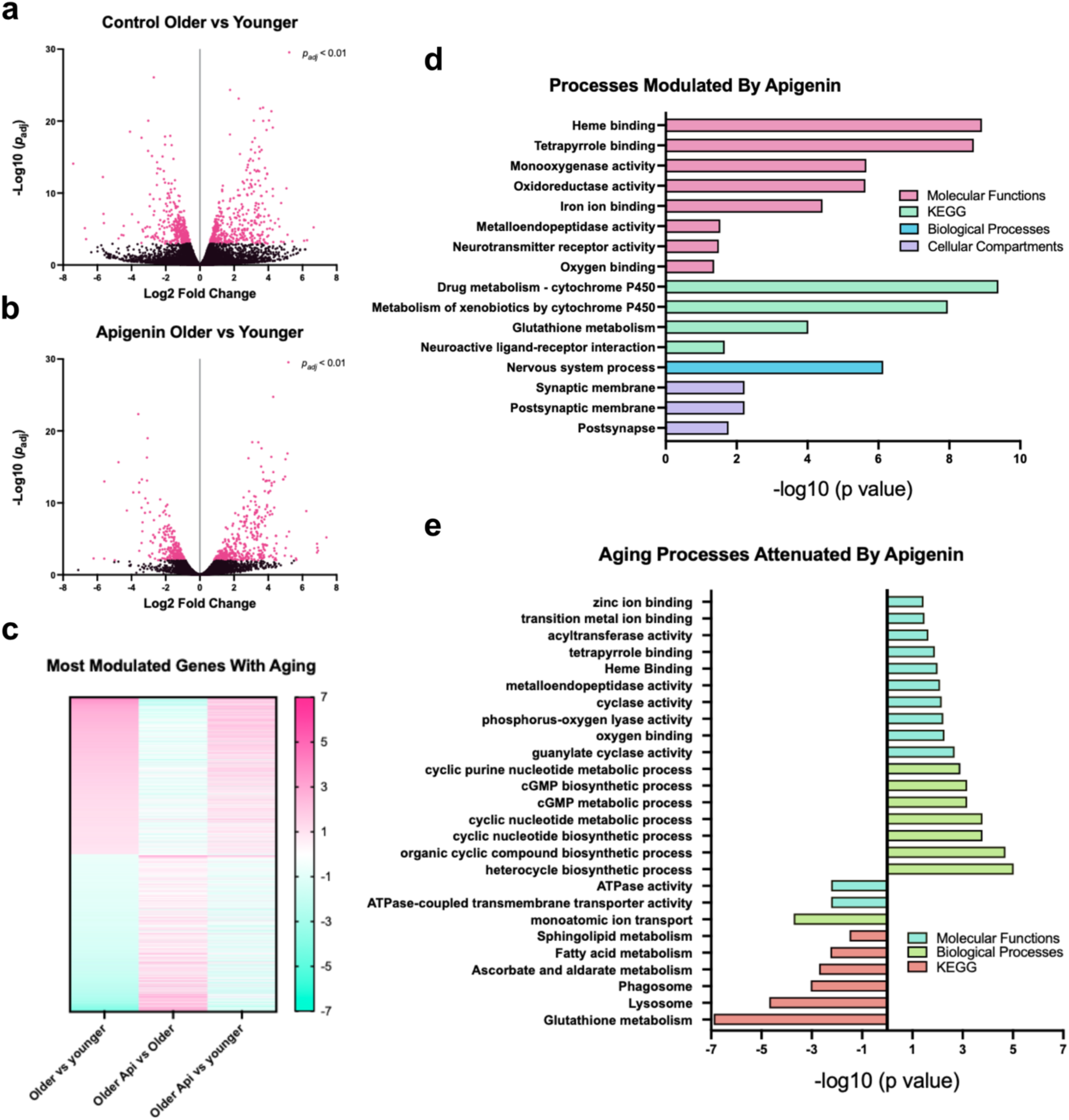
**Anti-aging transcriptomic effects of apigenin**. Volcano plot showing significant differentially expressed genes in **a)** older vs. younger untreated wild-type *C. elegans*; and **b)** older untreated vs. older apigenin-treated wild-type animals, with significantly different transcripts shown in pink (*p*_adj_ < 0.01); **c)** Heatmap showing relative expression differences of the top 1,000 most increased and top 1000 most decreased genes/transcripts with aging, and relative expression same genes/transcripts in treated vs. untreated older animals, as well as treated older vs. younger controls; **d)** Most modulated GO terms with treatment; **e)** Most modulated GO terms associated with aging and also attenuated with apigenin treatment

Therefore, to more broadly understand how these transcript expression differences reflected biological processes affected by aging and apigenin, we examined patterns among the top 1000 most increased and decreased genes/transcripts with aging. We found that apigenin broadly reversed transcriptional patterns among these transcripts in old animals (i.e., most transcripts that were increased with aging were reduced with apigenin, and vice versa; **Fig. 4c, middle panel**). Additionally, day 10 treated worms were more transcriptomically similar to day 5 untreated animals, as reflected by less pronounced “aging” gene increases/decreases (**Fig. 4c, right panel**). A GO analysis of the most increased and decreased differentially expressed genes between day 10 treated and day 10 untreated animals also revealed enrichment of oxidative stress-related metabolic processes in the nervous system (**Fig. 4d**), which was consistent with our data showing that apigenin may protect against age-related neurodegeneration and oxidative stress *in vivo*. Next, to identify which of apigenin’s transcriptome effects were most relevant to its anti-aging properties, we specifically examined genes/transcripts that were increased and decreased with aging but no longer increased or decreased with chronic apigenin treatment (i.e., transcripts specifically “reversed” in terms of their expression by apigenin). GO analyses of this list showed that processes that increased with aging but were reduced with apigenin converged on stress defense and homeostasis (redox buffering, lysosomal clearance, ion transport, and lipid metabolism), while processes that decreased with aging but were increased with apigenin centered around stress response and redox buffering (**Fig. 4e**). Together, these data support a model in which apigenin protects against age-associated transcriptome shifts by modulating biological pathways related to oxidative stress and redox homeostasis.

Oxidative stress is a well-established contributor to AD (59,60), and Aβ (a key pathological protein in AD) has been linked to oxidative stress (61). Given our data presented above, and previous reports that apigenin protects against age-related neurodegeneration and oxidative stress-associated pathway activation with aging (62–64), we hypothesized apigenin could also have protective effects against AD-relevant proteotoxicity. To test this idea, we performed lifespan analyses on transgenic *C. elegans* strains expressing pan-neuronal human Aβ_1-42_ and tau. We found that apigenin significantly increased median lifespan in Aβ-expressing worms (**Fig. 5a**), and analyses of pharyngeal pumping (**Fig. 5b**) and crawling velocity (**Fig. 5c**) at median lifespan (day 7) revealed significant enhancement of neuromuscular function as well. Additionally, fluorescence microscopy analyses showed that apigenin-treated Aβ-expressing animals had significantly reduced 447 nm autofluorescence, reflecting reduced accumulation of damaged/aggregated macromolecules, at the same time point (**Fig. 5d,e**). Together, these data suggest apigenin may have potential to modulate Aβ proteotoxicity.

**Fig. 5.**
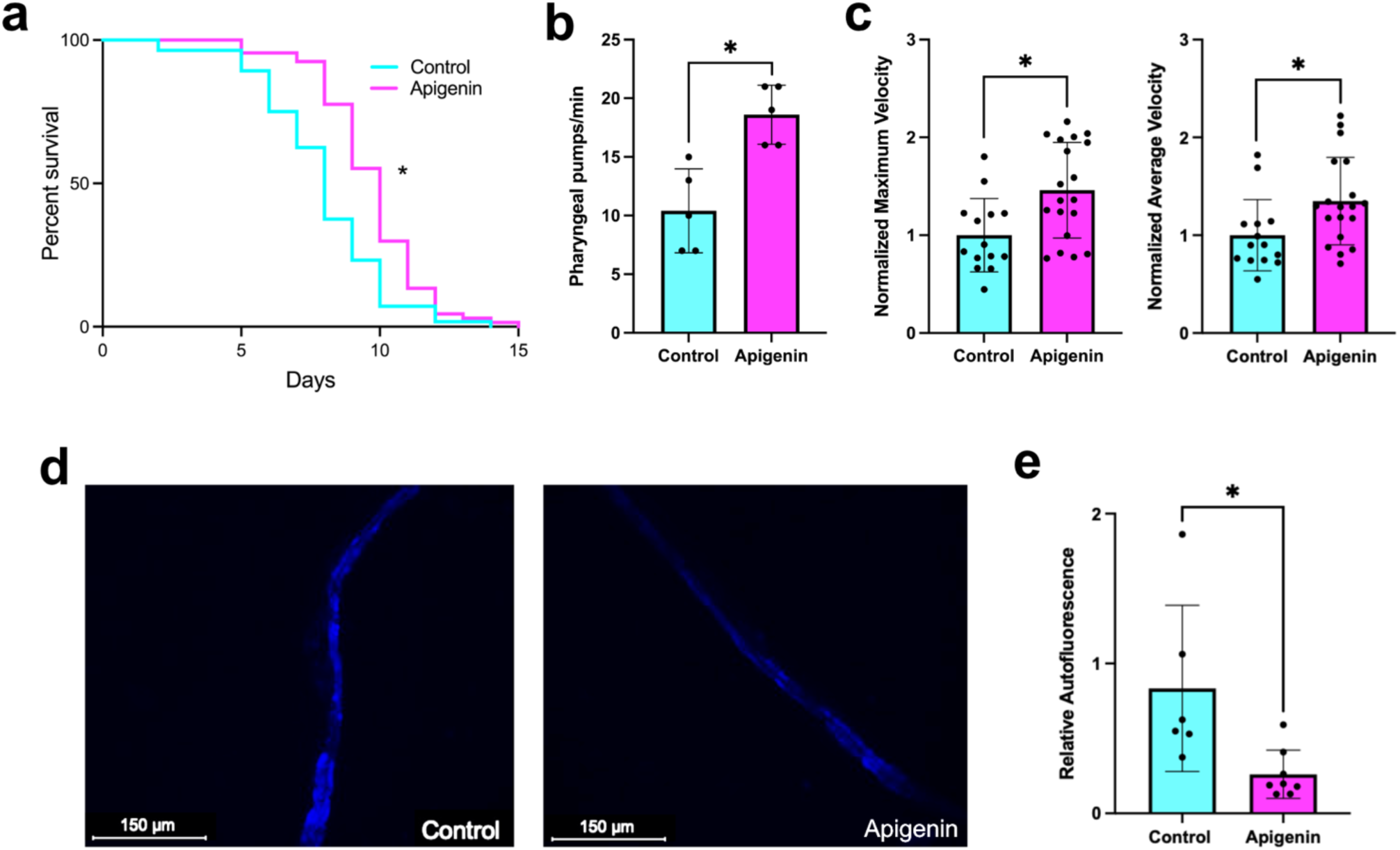
Apigenin improves healthspan in. **A**β**-expressing *C. elegans*. a)** Median lifespan was significantly extended in apigenin-treated worms (Control n = 56, Apigenin n = 67) expressing pan-neuronal Aβ. In the same animals, neuromuscular function was significantly enhanced by apigenin as reflected by increased **b)** pharyngeal pumping rates (*n* = 5/group) and **c)** average and maximum crawling velocity (*n* = 14-19/group). **d)** Representative age-related autofluorescence (λ = 447 nm) images. **e)** Quantification of age-related autofluorescence intensity. All measurements were assessed at day 7 (*n* = 6-8/group). \**p* < 0.05

The relationship between tau and oxidative stress is less established than that of Aβ. However, oxidative stress and tau hyperphosphorylation are thought to be part of a vicious cycle in which each promotes the other (65). To determine if apigenin’s protective effects might extend to tau-related pathology, we performed lifespan and healthspan analyses on control and apigenin-treated worms expressing pan-neuronal pathological human tau. Similar to our findings in Aβ transgenic animals, we observed a significant increase in median lifespan in apigenin-treated tau-expressing animals (**Fig. 6a**), accompanied by significant increases in neuromuscular function (**Fig. 6b,c**) and significant decreases in 447 nm autofluorescence (**Fig. 6d,e**). Together, these data indicate that apigenin may have broad protective effects against AD-associated proteotoxic stress in *C. elegans*.

**Fig. 6.**
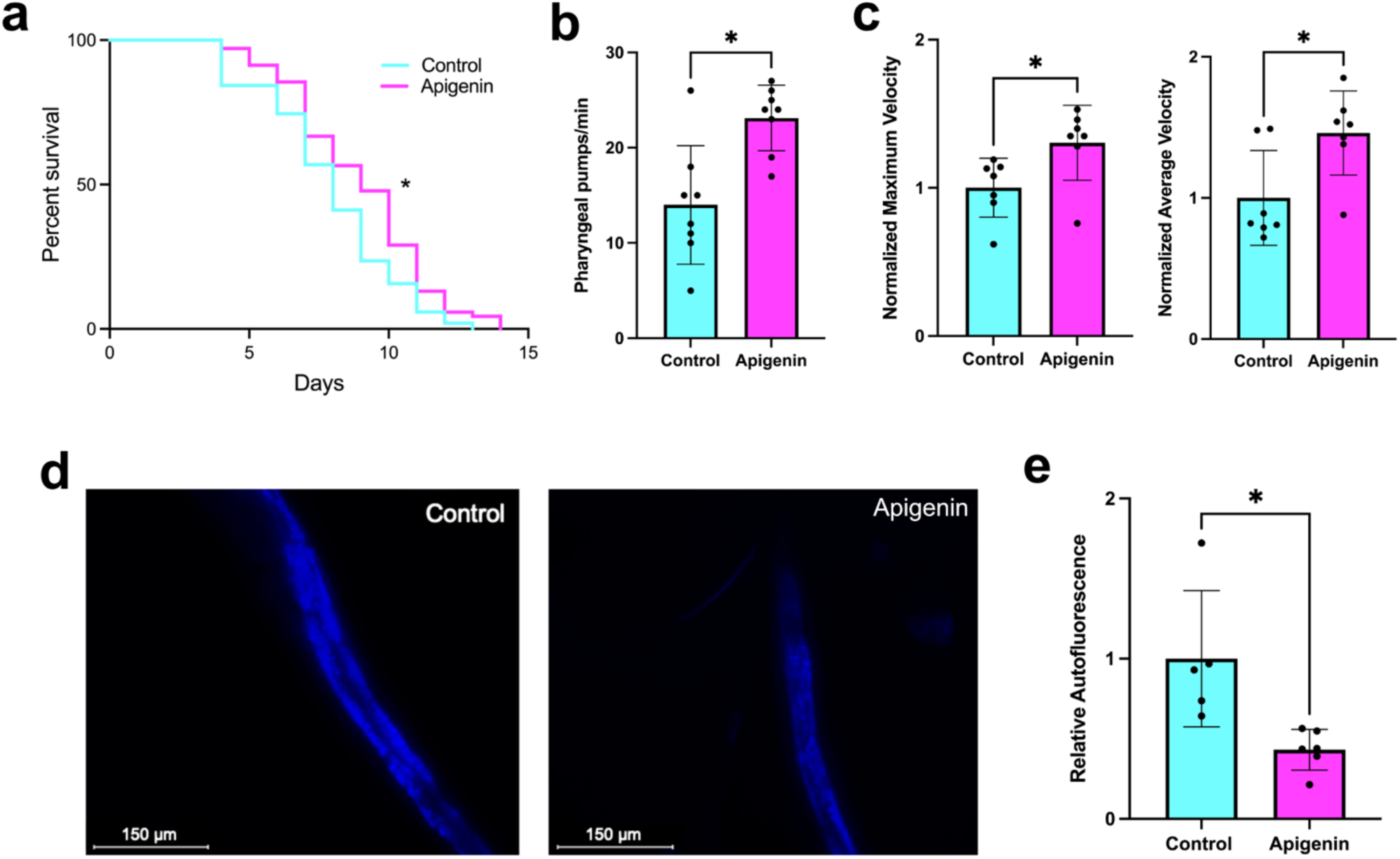
Apigenin improves healthspan in tau-expressing *C. elegans*. **a)** Median lifespan was significantly extended in apigenin-treated worms (Control n = 51, Apigenin n = 69) expressing pan-neuronal tau. In the same animals, neuromuscular function was significantly enhanced as reflected by increased **b)** pharyngeal pumping rates (*n* = 8/group) and **c)** average and maximum velocity (*n* = 7/group). **d)** Representative age-related autofluorescence (λ = 447 nm) images. **e)** Quantification of age-related autofluorescence intensity. All measurements were assessed at day 7 (*n* = 5-6/group). \**p* < 0.05

Finally, although we observed significant increases in median lifespan and healthspan in pan-neuronal Aβ and tau worms treated with apigenin, it remained unclear if apigenin specifically protects against individual pathological proteins, or if it simply attenuates accelerated aging effects due to proteotoxicity. Therefore, to better understand if apigenin was protective against Aβ and tau proteotoxicity outside of a purely neuronal context, we utilized *C. elegans* models expressing either human Aβ_1-42_ or pathological tau specifically in muscle. We found that apigenin significantly delayed paralysis onset in transgenic animals expressing temperature-inducible Aβ in muscle (**Fig. 7a**). Similarly, we observed a significant delay in locomotor decline of apigenin-treated worms with constitutive tau expression in muscle, as reflected by greater average and maximum velocities (**Fig. 7b**). These findings suggest that apigenin improves resistance to proteotoxic stress in general, with protective effects beyond neurodegeneration that may preserve physiological function across tissue types.

**Fig. 7.**
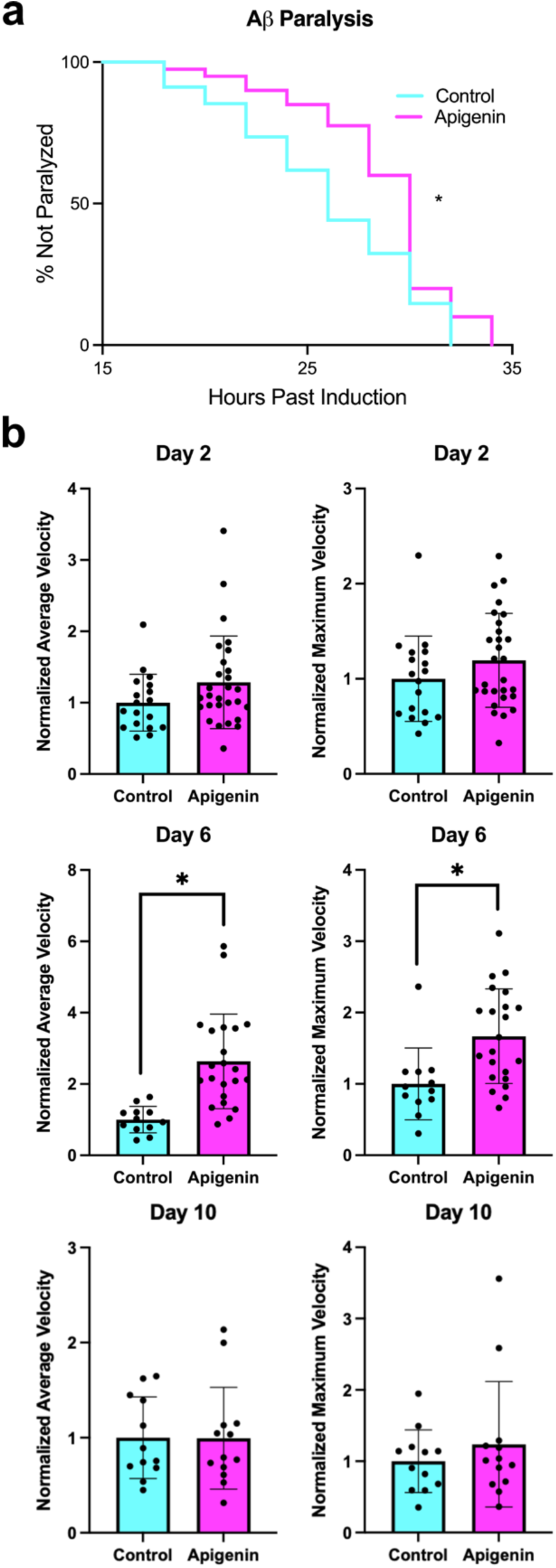
Apigenin protects against AD protein-induced paralysis. **a)** Aβ-induced paralysis (CL4176 strain) was significantly delayed with apigenin treatment. All worms were completely paralyzed by 35 hours post-induction (Control *n* = 34, Apigenin *n* = 40). **b)** Tau-induced locomotor decline was also delayed by apigenin treatment, as reflected by significantly increased average and maximum velocity in control vs. apigenin-treated tau-expressing animals (KAE112; *n* = 12-30/group). \**p* < 0.05

## DISCUSSION

Apigenin is a naturally occurring compound found in fruits, vegetables, and tea (66). It is among the most extensively studied and well-characterized dietary flavonoids, with reported therapeutic potential in multiple disease models (47). In fact, apigenin has been shown to exert beneficial effects in models of cancer (67–71), cardiovascular disease (72,73), diabetes (74,75), viral infection (76–78), kidney dysfunction (79,80), and multiple neurodegenerative diseases (81–84). Current literature suggests that the mechanisms underlying apigenin’s therapeutic effects may be context dependent. For example, in cancer models, multiple studies have shown that apigenin induces both cell cycle arrest (25,85) and p53-mediated apoptosis (86–89). Yet, in other disease models, data suggest apigenin’s beneficial effects primarily involve either antioxidant activity (86) or anti-inflammatory properties (11,90–93), with multiple studies documenting reductions in both (66,94,95).

Still, the degree to which apigenin is protective against aging and AD-related proteotoxicity specifically has not been thoroughly investigated. The key findings of the present study are that apigenin attenuates physiological aging and age-related functional declines in *C. elegans*, as well as neurodegeneration-relevant proteotoxicity *in vivo*. Importantly, unbiased transcriptome-wide profiling revealed that the most heavily impacted biological pathways in this context are associated with oxidative stress signaling and redox homeostasis, supporting apigenin’s potential as a functional dietary intervention to combat physiological aging.

Several recent studies have examined the therapeutic effects of apigenin and/or its metabolites on lifespan and aging phenotypes, yet reports have suggested various mechanisms with no clear consensus (96–98), To our knowledge, only two studies in higher organisms have tested the effects of apigenin on the aging brain, and although both demonstrated beneficial effects, one study suggested an antioxidant mechanism of action, while the other suggested primarily anti-inflammatory properties (11,99)—and none of these studies examined healthspan and neuromuscular health *per se*. These findings highlight a lack of consensus on the biological pathways by which apigenin influences aging and neurodegeneration. To address this, we performed a broad analysis of apigenin’s effects on aging in wild-type *C. elegans*, incorporating lifespan, functional, and transcriptomic readouts. First, we found that apigenin increases both median lifespan and neuromuscular function while reducing age-related autofluorescence (a marker of macromolecular damage accumulation) in wild-type worms, consistent with reports of apigenin’s effects on neuronal aging (11,99) and muscle physiology (100,101) in rodents. Additionally, we observed a significant age-associated accumulation of neuronal blebbing in control worms, which did not reach significance in apigenin-treated animals, similar to *in vitro* reports showing apigenin protects against peripheral nerve degeneration (64). We then tested apigenin’s effects in multiple transgenic *C. elegans* reporter strains at mid-life, with an emphasis on reporters for oxidative stress and inflammation pathways. We found that apigenin-treated animals had reduced expression of *daf-16*, a FoxO (a transcription factor acting as a master regulator of cellular homeostasis, longevity, and stress resistance (102)) ortholog and key mediator of stress response and longevity in *C. elegans*, suggesting lower age-associated cellular stress in treated animals (103). We also observed decreased expression of both *skn-1b/c* and *gcs-1p*, key indicators of oxidative stress signaling in *C. elegans*, with no changes in muscle mitochondria reporters, consistent with a mechanism of reduced oxidative stress rather than mitochondrial remodeling (104–106). We note, however, that we did not directly measure mitochondrial function or reactive oxygen species (ROS) production, and therefore could not distinguish between reduced upstream stress signaling and direct effects of apigenin on mitochondrial or redox processes. Importantly though, we found that apigenin reduced expression of *mpk-1* (the *C. elegans* ERK/MAPK ortholog) but not *nlp-29p* (a marker of innate immune activation). Given that ERK/MAPK signaling is a sensor of intracellular stress, the attenuation of age-related *mpk-1* increases in conjunction with unchanged *nlp-29p* expression suggest that apigenin targets core stress networks as opposed to generalized anti-inflammatory signaling pathways (107–110). Collectively, these reporter data indicate reduced activation of canonical stress response pathways during aging with apigenin treatment, consistent with a model in which apigenin may lower basal cellular stress.

Importantly, although hormetic mechanisms have been proposed to underlie the beneficial effects of many phytochemicals, including apigenin, the reduced expression of stress response reporters we observed in the present study does not support a chronic hormesis response (111,112). While we did not assess early or transient responses and therefore cannot completely exclude a role for hormesis, our findings suggest apigenin operates through alternative mechanisms that reduce basal age-associated stress signaling while preserving homeostatic systems (rather than inducing stress adaptation). Interestingly, apigenin was recently demonstrated to reduce ROS accumulation by preserving SIRT1 levels in a mouse model of heart injury, and computational analyses have predicted that apigenin may directly bind to and activate SIRT1 (62). This finding potentially explains the diverse therapeutic effects observed across studies, as SIRT1 is a deacetylase that regulates multiple key processes intersecting with lifespan, neurodegeneration, obesity, heart disease, and cancer (113). Both SIRT1 and the *C. elegans* ortholog *sir-2.1* modulate multiple biological processes associated with aging (114), including the regulation of FoxO/*daf-16* (115,116) signaling and oxidative stress responses (117,118). These literature insights closely align with our experimental observations too, in that apigenin may coordinate its multi-systemic benefits through *sir-2.1* upstream of *skn-1* and *daf-16*, although this remains to be tested directly. Future studies should examine the effects of apigenin in transgenic *sir-2.1* loss- and gain-of-function mutant animals to determine whether *sir-2.1* activity is necessary for its healthspan effects.

To provide broader insight into the effects of apigenin on aging biology, we performed transcriptomics (RNA-seq). We found that age-related transcriptional shifts were blunted in apigenin-treated worms, consistent with our autofluorescence and reporter strain imaging analyses. Treatment was also associated with a broad “reversal” of many age-dependent gene expression patterns in mid-life animals, driven primarily by genes reflecting oxidative stress-related metabolic processes in the nervous system (in line with our *in vivo* observations of reduced oxidative stress and preserved neuron integrity in treated animals). Additionally, some pathways/transcriptional signatures were “maintained” by apigenin during aging (i.e., declined in untreated aging worms but were preserved with treatment). These were enriched for genes/transcripts involved in lysosomal and phagosomal pathways, ATPase-coupled transporters, ion homeostasis, redox metabolism, and lipid metabolic processes. Key antioxidant buffering systems (glutathione and ascorbate/aldarate metabolism (119,120)) also declined with age but were maintained with apigenin treatment, consistent with the idea of preserved redox homeostasis. Notably, *C. elegans* often upregulate a suite of oxygen- and heme-binding enzymes as part of an age-associated oxidative stress response (121,122), and in our data, we observed an enrichment for cyclic-nucleotide signaling, heme-binding redox enzymes, and metalloproteases among genes that were increased with aging but attenuated by apigenin. These transcriptomic analyses indicate apigenin suppresses redox-sensitive stress signaling networks in aging *C. elegans*, while preserving some key cellular homeostatic processes that decline with age. Interestingly, this dual therapeutic action mirrors another naturally occurring dietary bioactive compound, α-ketobutyrate, which has specifically been shown to interact with *sir-2.1* (extending lifespan and healthspan mediated by enhanced autophagy and redox-buffering) (123).

Finally, because oxidative stress is a well-characterized feature of AD (124) and apigenin has shown promise as a therapeutic in models of neurodegenerative disease (125), we evaluated the effects of apigenin in transgenic *C. elegans* strains expressing AD-associated pathological proteins (i.e., Aβ and tau). Although *C. elegans* has a simple nervous system and does not develop AD-like neurodegeneration (limiting its ability to recapitulate human disease), it has many conserved molecular stress response pathways and can serve as a cost-effective platform for testing compounds that may protect against toxicity associated with disease-relevant proteins (126,127). In animals expressing either Aβ or tau in neurons, apigenin consistently exerted protective effects on lifespan, neuromuscular function, and macromolecular damage accumulation. This was consistent with previous studies on apigenin’s protective effects against AD-pathology in mice (81) and in cultured neurons (82,128). However, we also found that for *C. elegans* strains expressing either human Aβ or tau in muscle, apigenin significantly delayed paralysis, suggesting that its protective effects extend beyond neuronal disease-associated proteotoxicity, with the potential to preserve physiological function across multiple tissue types.

Collectively, the data from this study lend additional support to the idea that apigenin may be a promising anti-aging/neurodegeneration therapeutic. We also provide new mechanistic insight into its therapeutic effects, showing that apigenin acts to suppress the age-related activation of redox-sensitive stress response signaling while preserving homeostatic processes that deteriorate with age and disease. Key limitations to our study are that the reporter strain and RNA-seq data were cross-sectional and not based on cause-and-effect evidence. Additionally, *C. elegans* has inherent limitations when modeling neurodegeneration as they lack a centralized brain structure and true glial cells like vertebrates. Based on our findings, future studies dissecting apigenin’s mechanisms of action should test its effects in transgenic *C. elegans* strains with mutations in *sir-2.1*, *skn-1*, and *daf-16* while making more direct measures of proteostasis and oxidative stress. Additional studies will also be needed to determine how similar pathways are modulated with aging and apigenin in higher model organisms to establish translational relevance. Ultimately, these data add to the literature suggesting apigenin as a promising intervention for age-related functional declines and identify stress responsive/homeostatic pathways that may underlie its effects.

## Data Availability Statement

All raw RNA sequencing data generated in this study will be made publicly available on the Gene Expression Omnibus (GEO) server at the time of publication.

## Author Contributions

**R.A.G.** designed the study, wrote the paper, performed lifespan and behavioral analyses, generated microscopy data, generated/analyzed RNA sequencing data, and performed paralysis assays. **L.H.E.P.** assisted with lifespan assays and behavioral analyses, and with manuscript drafting. **D.W.** assisted with lifespan assays and manuscript editing support. **T.J.L.** designed the study, wrote/edited the paper, assisted with all *in vivo* and transcriptome analyses, provided conceptual insight, and provided funding.

